# Nuclear Packing Sets Fluidity Along the Epithelial to Mesenchymal Spectrum

**DOI:** 10.1101/2025.10.29.684532

**Authors:** Karen Yu, Alexander J. Devanny, Laura J. Kaufman

## Abstract

Investigations of jamming in cancer have produced numerous phase diagrams intending to map fluidity across the epithelial-to-mesenchymal transition (EMT). Here, we use coalescence of homotypic and heterotypic multicellular spheroids to examine and carefully probe these phase spaces. Small changes in cellular EMT status result in full traversal of the solid-to-fluid continuum. We propose that stiff nuclei impede cell motion and spheroid coalescence and find that by softening nuclei, fluidization of an otherwise solid-like system occurs. Changes in fluidity during coalescence is fully captured by static cellular properties, such as internuclear spacing and nuclear shape, that can be assessed in individual non-interacting spheroids. We combine these quantities into an effective nuclear packing metric that depends on nuclear occupancy and nuclear elongation. Together, these findings reveal that nuclear morphology and packing act as important indicators and determinants of fluidity.

## I. INTRODUCTION

Cell migration is a key component of processes such as embryonic development, wound healing, and metastasis. It has been proposed that these processes, as well as cell migration generally, can be viewed through the framework of jamming transitions. For example, cells in solid-like tumors unjam and migrate during metastasis, bronchial epithelia compress and flow in asthma, and drosophila development depends on a fluid-like transition during convergent extension [1–4].

Traditionally, jamming has been used to describe passive physical systems such as grains of sand [5]. The phase space that has been used to map jammed systems includes contributing factors of temperature, density, and load [6]. Such a jamming phase diagram can be mapped to biological systems, with active fluctuations, volume fraction, and supracellular stress as the relevant analogous parameters [7–9]. With the surge of studies of biological jamming in different contexts, other phase spaces have been proposed using parameters such as cell-cell adhesion, cell contractility, cell shape, and cell area [10–13]. These phase spaces have been mapped extensively with computational models [1,14–16], as well as with histological samples [10]. However, few studies have experimentally explored these proposed phase spaces, in part because tools are lacking to perturb these cellular features independently.

Of the many possible jamming parameters, cell shape has been shown in both experiments and simulations to have an important role in fluidity. Specifically, a jamming transition is observed as cells become rounder [2,17]. Beyond this, nuclear shapes and deformability have recently gained attention as playing a major role in mobility [11,18–24]. It has been reported that mobile cells have more elongated nuclei, as to navigate a crowded environment cells need to pass their nuclei between the surrounding cells or extracellular structures [25]. This leads to the idea that in tissues nuclear packing may play a role in mobility. Previously confluent cellular systems were understood to have a packing density of one, as there is little to no intercellular space; however, given cell capacity to deform, there has been growing recognition that cellular packing density may not be the most relevant parameter. Indeed, recently, it has been shown that the area fraction of the nucleus to the cytoplasm plays a role in mobility, with higher nuclear area fractions leading to denser packing and greater tissue stability [20]. However, there remains limited work that directly probes the nucleus in cellular systems either in computational studies or experimentally.

In this work, we use multicellular spheroid coalescence to probe the fluidity of different cell types and elucidate the role of the nucleus in jamming. Spheroid fusion has been used extensively to investigate cellular jamming, with macroscopic readouts such as the degree of fusion giving insight into cell motility and mechanical properties such as surface tension [11,26–31]. We find that small changes in epithelial-to-mesenchymal transition (EMT) status can recapitulate the full solid-to-fluid range of coalescence. We show that both elongated cell and nuclear shapes and loose nuclear packing are correlated with higher degrees of fluidity. When stiff nuclei are tightly packed, they are an impediment to fluidity - thus by softening the nucleus a formerly solid-like system can be fluidized. Furthermore, we show that heterotypic spheroids can be used to actively tune the fluidity of the system, allowing for careful investigation of jamming phase spaces. Our results confirm the finding that cell and nuclear shapes determine the onset of fluidity of our system with cell area having a small, independent additional contribution to the degree of fusion. Beyond this, we propose a metric that captures nuclear packing and shape, providing a single parameter structural predictor of fluidity.

## II. RESULTS

### A. Small Differences in EMT Status Are Sufficient to Drive a Solid to Fluid-like Transition

To probe the mechanisms and significance of fluid-solid behavior in cancer progression, we employ an isogenic cell line pair consisting of MCF10A (10A) and MCF10DCIS.com (DCIS), a model normal breast epithelial cell line and a minimally transformed 10A derivative, respectively. In brief, DCIS cells are derived from a xenograft of pre-malignant MCF10AT cells injected into immunodeficient mice such that ductal carcinoma in situ lesions form [22]. In this way, DCIS cells represent an early, non-invasive breast cancer that may eventually progress into invasive ductal carcinoma. 10A and DCIS cells share many characteristics including similar expression levels of epithelial and mesenchymal markers such as E-cadherin, N-cadherin and vimentin [33]. As a result, both cell lines are relatively epithelial in character. This experimental system is in contrast to previous works that examine differences between cell lines on extreme ends of the EMT spectrum [11].

To interrogate the fluidity or solidity of each cell type, we utilized a multicellular spheroid tumor model, which has seen use in studies of cancer progression and tissue dynamics [11,27–30,34–36]. Here, thousands of cells self-assemble into a microtissue roughly 200-300μm in diameter. To assess the fluidity of a given cell type, two spheroids are placed side by side and allowed to fuse into a single aggregate. In some cases, the two spheroids coalesce completely into a singular circular aggregate (**Fig 1(a), top**). Here, surface area is minimized in an apparently surface tension driven process, as in Newtonian fluids; thus, the cells are presumed to be in a fluid-like state. Alternatively, some spheroids stall during fusion (**Fig 1(b), bottom**). These systems exhibit behavior clearly deviating from liquid droplet fusion and are presumed to be solid-like in character. Qualitatively, this fluid-like vs. solid-like behavior can be easily distinguished by examining the end state of spheroid fusion experiments (**Fig 1(b)**). Stalled fusions have an elongated geometry, and therefore larger aspect ratio (AR), while completed fusions are nearly perfectly circular and have an AR close to 1. To quantitatively distinguish between these cases, we defined AR_fusion_ ≤ 1.2 as the threshold for a circular geometry. When assessing fluidity of 10A and DCIS cells, we primarily considered the case in which cell division was absent by treating cells with a proliferation inhibitor prior to spheroid formation (see **Methods**), as proliferation may act to fluidize otherwise solid-like systems [37]. Non-proliferative 10A spheroids begin to fuse but eventually stall at AR_10A_ = 1.63 ± 0.07, while DCIS spheroids quickly and completely fuse to an aspect ratio of AR_DCIS_ = 1.11 ± 0.05 (**Fig 1(b)**). Surprisingly, despite the minimal transformation and non-invasive character of DCIS [33], we observe a dramatic increase in tissue fluidity relative to 10A cells. This demonstrates that a minor change in the EMT status, causing no change in invasiveness, is capable of fluidizing an otherwise solid-like tissue.

**FIG 1.**
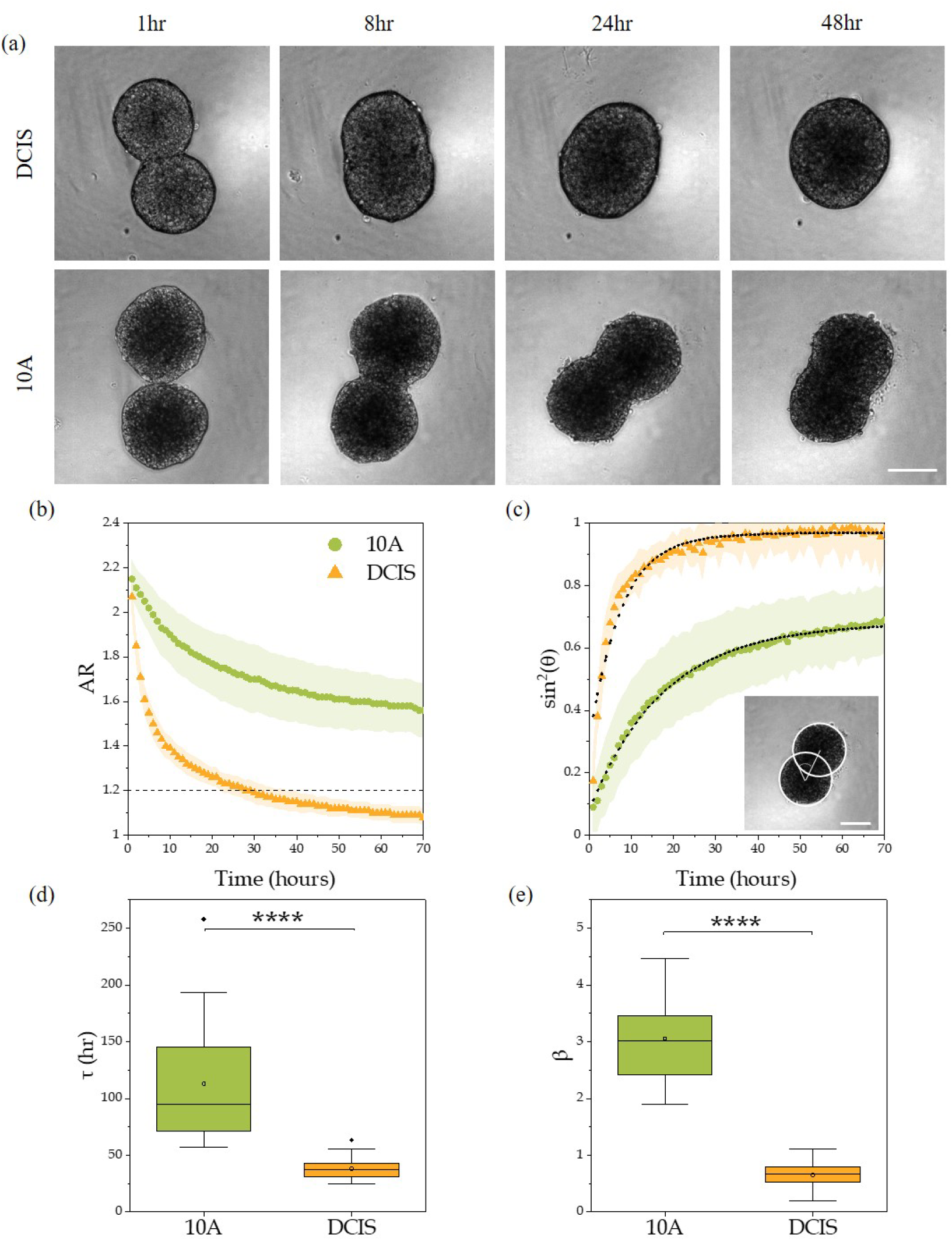
(a) Top: MCF10DCIS.com (DCIS) and bottom: MCF10A (10A) spheroids during 1-48 hours of coalescence. Scale bar is 200 µm. (b) Aspect ratio (AR) of the fusing 10A (green circles) and DCIS (orange triangles) spheroids over the duration of the experiment (N > 30 spheroids). Dotted line indicates the aspect ratio cutoff of a liquid-like system, AR ≤ 1.2. The AR ≤ 1.2 cutoff was determined as point at which coalescing spheroids were visually indiscernible from a single sphere. (c) The evolution of the angle of fusion via sin^2^θ, with θ the angle of fusion between the two spheroids as shown in the inset (N > 30 spheroids). Fits of the averaged data using the Kelvin-Voigt model (see Methods) are shown by the dotted line. Standard deviations are shown as shaded regions. Scale bar is 200 µm. (d,e) Box and whisker plots displaying τ and β obtained from fits of sin^2^θ for individual sets of fusing spheroids (N ≥ 30). Whiskers display range of data excluding outliers. Both τ (p_MW_ < 0.0001) and β (p_t-test_ < 0.0001) are significantly different between 10A and DCIS spheroids. Legend in (b) applies to (b-e).

A variety of modelling approaches exist for quantifying fusion dynamics [11,27,29–31,35,38–40]. In the simplest and most popular model, tissues are modeled as passive Newtonian fluids. However, modeling fusion as viscous fluids driven by surface tension fails to describe stalled coalescence [11]. Further, many liquid drop models use metrics that consider the neck length (the contact length between the two spheroids), but such models lack the ability to discern the difference between neck growth and meaningful increases in fusion. Given these issues, it is best to use a model that tracks the angle of fusion and can account for stalled coalescence.

Here, we treat the spheroids as homogenous incompressible Kelvin-Voigt materials and track the angle of fusion over time (**Fig 1(c))**. The Kelvin-Voigt model is the simplest model that has been successfully used to describe arrested coalescence in similar systems [30]. This fitting method considers spheroid fusion as driven by effective shear viscosity, shear modulus, and surface tension (see **Methods**). From fits, a fusion timescale, τ, and degree of fusion, β, are obtained, with β reflecting the aspect ratio of post-fusion spheroids (**Appendix A: Fig 7**) [30]. We see that the aspect ratio cutoff of AR_fusion_ ≤ 1.2, which denotes fluidity, corresponds to β_fusion_ ≤ 1.2. Both τ and β show significant differences when comparing non-proliferative 10A and DCIS spheroids (**Fig 1(d-e)**). Characteristic of a jammed system, the 10A spheroids exhibit long relaxation timescales (τ_10A_ = 113 ± 49 hrs), much longer than the duration of the experiment. In contrast, the timescale obtained for DCIS spheroids (τ_DCIS_ = 38 ± 9 hrs) reflects the significantly faster fusion progression. The extracted degrees of fusion also decrease from β_10A_ = 3.1± 0.7 to β_DCIS_ = 0.7 ± 0.2, reflecting their stalled and fully coalesced states, respectively. Although we focus primarily on non-proliferative spheroids in this work, we note that in proliferative spheroids, an increase in fusion ability is observed for 10A spheroids while little change in DCIS spheroid fusion is seen (**Appendix B: Fig 8**). This suggests that in a solid-like system, perturbations from proliferation appear to aid in fusion.

### B. Cell Shapes

To better understand the transition between fluid and solid behavior, we probed the internal structure of spheroids using the cell shape index, 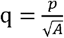, where *p* and *A* are the perimeter and area of the cell, respectively. The shape index serves as an indicator of tissue fluidity and allows one to infer dynamic behaviors from static structure [3,17,41–43]. Specifically, elongated and therefore higher shape index cells have been found to be more mobile than round cells. To this end, we fixed and stained individual spheroids 24 hours after initiating formation. Spheroids were then imaged and segmented to obtain shapes (see **Methods**).

Qualitative examination of fixed individual spheroids reveals that DCIS spheroids have more disordered cell packing and are larger than 10A spheroids (**Fig 2(a)**). Further examination of the internal structure reveals that both 10A and DCIS spheroids are composed of elongated cells at the interface with the medium and more compact cells within the spheroid bulk, with q_10A,edge_ = 4.79 ± 0.59 and q_DCIS,edge_ = 5.21 ± 0.79 (**Fig 2(b)**) corresponding to highly elongated cell geometry in both cases. We propose that this elongated outer rim of cells observed across both 10A and DCIS individual spheroids facilitates the early relatively rapid fusion when either 10A or DCIS spheroids are placed in near vicinity and come into contact initially. However, as time passes and fusion progresses, the cells initially at the periphery become less elongated and rounder in shape, reflecting a transition from surface to bulk geometry (**Appendix C: Fig 9**).

**FIG 2.**
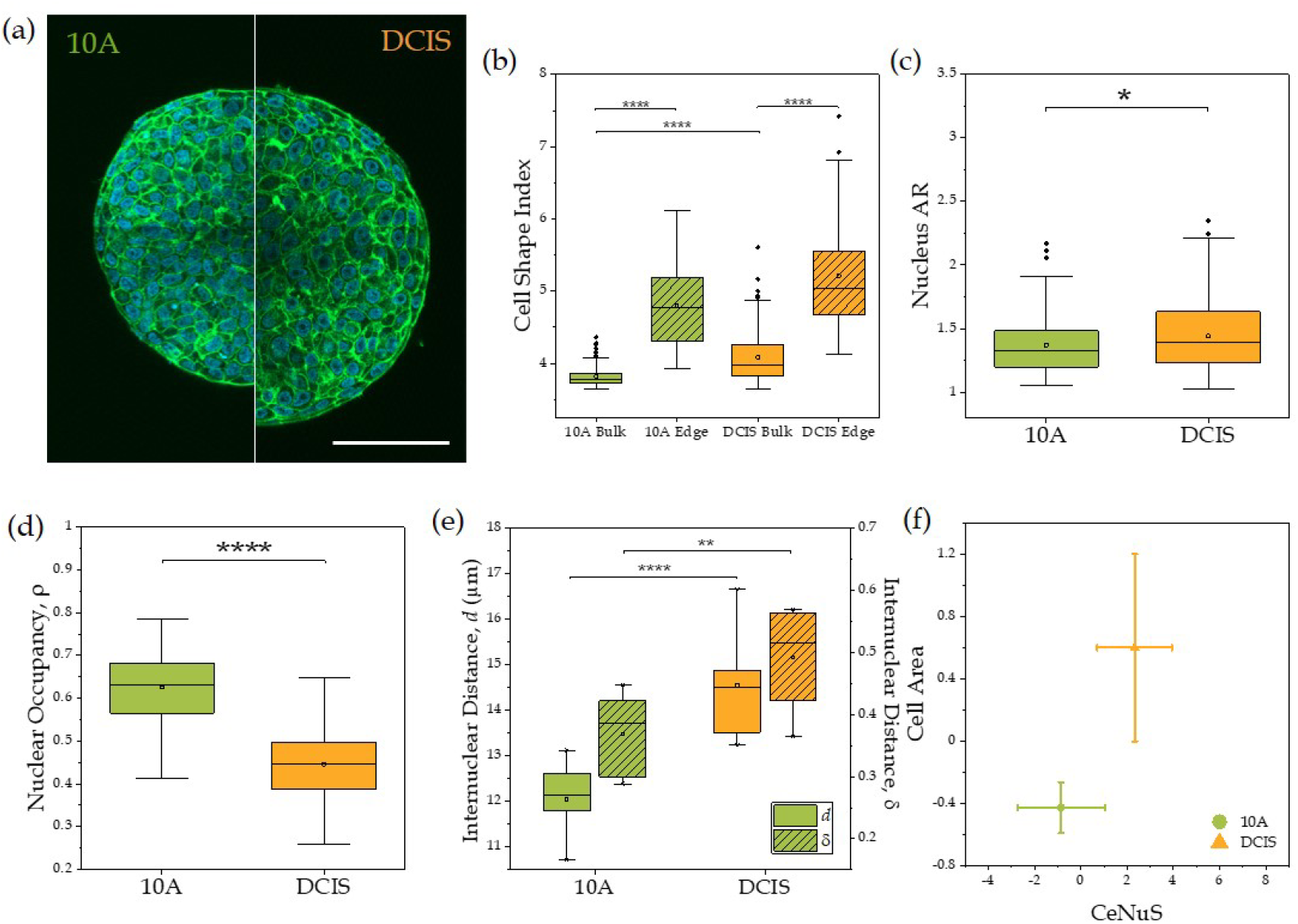
(a) Actin (green) and DAPI (blue) stains of individual spheroids. Left shows a 10A spheroid and right is a DCIS spheroid. Scale bar is 100 µm. (b) Cell shape index of 10A and DCIS cells in individual spheroids both in the bulk and at the periphery of the spheroids (N > 40 cells). (c) Nuclear aspect ratio (AR) (N > 100 cells) is significantly different between 10A and DCIS spheroids (p_MW_ < 0.05). (d) Nuclear occupancy (N > 40 cells) is significantly higher in 10A cells compared to DCIS cells in individual spheroids (p_t-test_ < 0.05). (e) Internuclear distance in absolute distance, *d*, (p_t-test_ < 0.0001) and normalized by nuclear size, *δ*, (p_t-test_ < 0.01) for 10A and DCIS cells in spheroids (N ≥ 3 spheroids for each condition). For all box and whisker plots, whiskers display range of data excluding outliers. (f) 10A and DCIS cells plotted in the Cell Area vs CeNuS phase space (N ≥ 6 spheroids for each condition). Error bars display standard deviations.

The interior bulk cells of the 10A and DCIS spheroids exhibit differences in shape index that are greater than that of edge cells and correlate more fully with observed differences in fusion behavior. 10A bulk cells appear as relatively regular polygons with a shape index of q_10A_ = 3.82 ± 0.15. This value corresponds closely with the shape index of 3.81 that has been identified in simulated monolayers as a shape threshold denoting the boundary between fluid and jammed solid phases [17]. This 10A shape index is consistent with the limited fusion capabilities of the 10A spheroids. DCIS spheroids, on the other hand, contain significantly more elongated cells, with a mean shape index of q_DCIS_ = 4.08 ± 0.35. In sum, the studied system confirms the previously identified link between cell shape and fluidity.

### C. Nuclear Contributions

Spheroids are densely packed microtissues with little to no empty space between cells and therefore have a packing fraction near unity. Additionally, the nucleus is known to be one of the stiffest and largest compartments of the cell [44]. As such, in order for a cell to navigate a crowded environment, such as the interior of a spheroid, the nucleus must be squeezed between surrounding cells [45]. Previous work has also shown that elongated mobile cells have elongated nuclei [11], indicating that nuclear shape could be another useful and perhaps more direct indicator of fluidity beyond cell shape.

We hypothesized two means through which the motility of cells in our system might be impacted by nuclei. First, we hypothesized that similar to cell shapes, differences in nuclear shape would be observed between fluid-like and solid-like cells. Given that nuclei are more ellipsoidal and have less tortuous shapes than cells, for nuclei we measured aspect ratio as opposed to shape index, in accordance with prior studies [10]. Consistent with previous work, we find that the more mobile and elongated DCIS cells have larger nuclear aspect ratios (**Fig 2(c)**). Second, barriers to cell motility can be reduced by reducing the packing fraction of the nuclei. Examining the difference between 10A and DCIS cells, we see that the nucleus occupies the majority of the cross-sectional cell area in 10A cells, ρ_10A_ = 0.63 ± 0.09, whereas for DCIS cells the nucleus occupies a significantly smaller proportion of the available space, with a nuclear fraction of ρ_DCIS_ = 0.45 ± 0.08 (**Fig 2(d)**), suggesting a lower nuclear packing density in DCIS. This effect results largely from differences in cell area, which are clearly evident by examining the differences between spheroid sizes (**Fig 2(a)**).

To directly probe the nuclear packing density, we measured the absolute internuclear distance, *d*, by averaging the 6 nearest neighbor internuclear distances of each cell within each spheroid (**Fig 2(e)**). In accordance with similar work [20], we then normalized the absolute internuclear distance to provide a system-independent assessment of nuclear packing (**Fig 2(e)**). In brief, the absolute internuclear distance was normalized to the average minor axis length of the ellipsoidal nuclei for that specific spheroid,*l* _nuc_, to obtain the normalized internuclear spacing, *δ* :

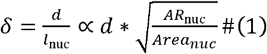

where *l*_nuc_ indicates the characteristic minor axis diameter of the nucleus for a given spheroid, which is a function of the aspect ratio, *AR*_*nuc*_ and area of the nuclei, *Area*_nuc_. It is important to recognize that the definition of the characteristic nuclear diameter as the minor axis of the fitted ellipse reveals that the effective nuclear packing can be decreased either by directly reducing the space occupied by the nucleus or by elongating the nucleus. Thus, this normalized quantity depends on both the nuclear area and shape.

When examining both the absolute and normalized internuclear distance, we find that the round 10A nuclei are densely packed, with typical nuclear spacing much less than the diameter of a nucleus. In contrast, DCIS nuclei are more loosely packed, with higher absolute and normalized internuclear spacing, which is a result of both lower nuclear area occupancy and more elongated nuclei (**Fig 2(e)**). The DCIS cells may thus be able to move more easily, without having to further squeeze their nuclei between other nuclei.

To encapsulate the simultaneous influence of changes in cell shape, nuclear shape, and nuclear packing density, we turn to the shape metric proposed by Gottheil et al. [10]. This metric, CeNuS, which combines cell and nuclear shape, is defined as

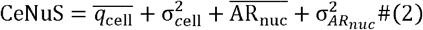

and captures the structural information encoded in both cellular and nuclear shape distributions. Here,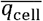 and 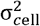 are the standardized median cell shape index and the corresponding variance, respectively and,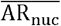 and 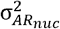 are the standardized median nucleus aspect ratio and the corresponding variance, respectively. This aggregate metric, when plotted against the projected cellular area (which reflects cell volume) [46], maps out a phase space that determines the fluid-like or solid like behavior of the tissue. More fluid-like tissues have a larger CeNuS value and/or larger cell area. Following the method described previously [10], we computed these parameters for individual 10A and DCIS spheroids (see **Methods**). We note that with our internal normalization, CeNuS values span a range of −6 to 6, a comparable range to previous studies, whereas our standardized cell area range is slightly smaller [10].Independent of normalization metrics, 10A and DCIS spheroids are well-separated in the Cell Area vs CeNuS phase space (**Fig 2(f)**). DCIS notably display an increased cell area and CeNuS, both of which suggest enhanced fluidity.

Based on this area-shape phase space and our observations regarding nuclear packing, we propose that the nucleus is the bottleneck in 10A fusion, and 10A cells are unable to rearrange due to the dense packing of their stiff round nuclei. While nuclei have been discussed as a potential barrier to cell mobility in confined spaces [40], the effects of nuclear stiffness have not been directly probed to our knowledge. Thus, we attempted to soften the nucleus by treating the cells with Trichostatin-A (TSA), an inhibitor of histone deacetylases [48]. As a result of softening the nucleus, the previously solid-like 10A spheroids now fuse significantly more and faster (**Fig 3(a)**). When treated with TSA, τ and β both decrease from a τ of 109 ± 53 hrs to 49 ± 13 hrs and β of 3.3 ± 0.9 to 1.4 ± 0.6 (**Fig 3(b-c))**.

**FIG 3.**
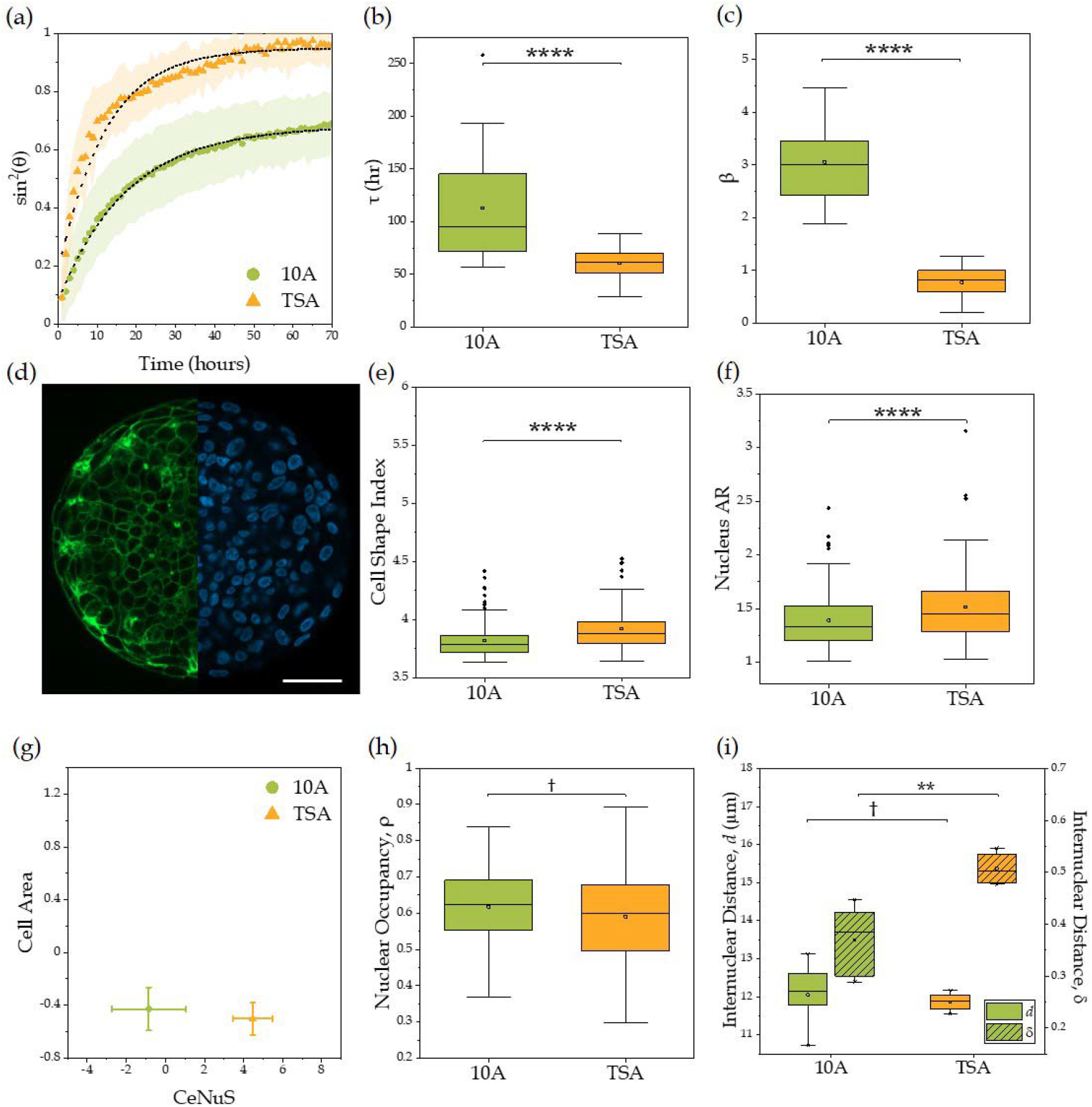
(a) TSA-treated 10A spheroids (orange triangles) fuse more than control 10A spheroids (green circles). (b,c) TSA-treated 10A spheroids (orange) fuse faster and more, as demonstrated by lower τ (p_MW_ < 0.0001) and β (p_t-test_ < 0.0001) values (N ≥ 16 spheroids). (d) Actin (green) and DAPI (blue) stains of an individual 10A TSA-treated spheroid. Scale bar is 50μm. (e) Cell shape index and (f) nuclear AR in 10A spheroids with and without TSA treatment (p_MW_ < 0.0001, N > 50 cells). (g) Cell area vs. CeNuS plot reveals that cell and nuclear shape change with TSA treatment but cell area does not, such that this treatment moves cells along the CeNuS axis only (N ≥ 6 spheroids). Error bars display standard deviations. (h) Nuclear occupancy, ρ, remains the same after TSA treatment (p_t-test_ < 0.0001, N > 100 cells). (i) Internuclear distance in absolute, *d*, (p_t-test_ > 0.05) and normalized distance, *δ* (p_t-test_ < 0.01, N ≥ 4 spheroids). Due to the elongated TSA nuclei, the normalized distance increases compared to control cells even though the nuclei remain the same size. For all box and whisker plots, whiskers display range of data excluding outliers.

Along with increased fusion ability in the 10A spheroids, both the cells and their nuclei become more elongated upon TSA treatment as can be seen via investigation of cells in individual spheroids (**Fig 3(d-f)**). In the CeNuS phase space, TSA-treated 10A spheroids thus shift horizontally along the CeNuS axis, while there is little change on the cell area axis (**Fig 3(g)**). As a result of the conserved cell and nuclear area, the nuclear fraction, and by extension the absolute nuclear spacing, are also unchanged by TSA treatment (**Fig 3(h-i))**. However, once nuclear spacing is normalized by nuclear diameter, an increase in effective nuclear spacing for TSA-treated spheroids is apparent (**Fig 3(i)**). This difference in spacing is dominated by changes in nuclear shape, as there is no change in nuclear (or cell) area. Thus, we see that though nuclear area fraction can be used as a proxy for absolute internuclear distances, nuclear shape must also be considered to accurately obtain nuclear spacing. For TSA-treated spheroids, the elongated nuclear shapes result in an increase in effective internuclear spacing, which increases fusion capability.

In total, this suggests that even when nuclei are densely packed in 10A spheroids, which would normally prevent fusion, softening of the nuclei with TSA and the resulting changes in cell and nuclear shapes allow for fluidization and therefore increased fusion. The increase in nuclear aspect ratio is a direct consequence of softer nuclei that more easily deform, presumably allowing cell-cell rearrangements. In contrast to 10A spheroids treated with TSA, TSA-treated DCIS spheroids remain relatively unchanged – they remain fluid, though fuse slightly faster, and there is little change in their shapes (**Appendix D: Fig 10**). DCIS spheroids are both more loosely packed and do not require further nuclear softening in order to fuse.

### D. Heterotypic Spheroids Allow for Traversal of Fluidity Phase Space

#### 1. Heterotypic 10A-DCIS spheroids

By employing the 10A-DCIS cell line system in combination with nuclear softening treatments, we demonstrate that the CeNuS-Area parameter space can be traversed, toggling between fluid-like and solid-like phases. For a more fine-grained map of this phase space, we next employed heterotypic spheroids composed of mixtures of fluid-like and solid-like cells. Indeed, physiological systems, in particular cancerous tumors, are highly heterogeneous [26,49,50], and characterizing how the proportions of different cell types within tissues translate to bulk tissue dynamics is of significant interest. Prior computational and experimental work in heterotypic cellular monolayers has demonstrated that altering heterogeneity can tune the jamming properties of the system [51,52].

We first probed the fusion capacity of a mixed spheroid system containing varying proportions of 10A and DCIS cells (**Fig 4(a))**. Adding DCIS cells to a predominantly 10A spheroid leads to a gradual change in fusion ability (**Fig 4(b)**). Even a small proportion of DCIS cells caused the spheroids to fuse more, as spheroids with just 10% DCIS cells had dramatically lower τ and β values than homotypic 10A spheroids (**Fig 4(c-d**)). When the spheroid was composed of at least 50% DCIS cells, the spheroids completely fused, reflected in τ and β values converging to the values observed in fusion of homotypic DCIS spheroids. Detailed examination of the CeNuS-Area phase space reveals that heterotypic spheroids display a gradual and relatively linear increase in both CeNuS and cellular area as the proportion of DCIS cells increases (**Fig 4(e)**), approaching the values observed in homotypic DCIS spheroids. Looking more closely, we find that the bulk shape index is not simply an average of varying proportions of two distinct cell shape populations. Instead, DCIS cells become more round as the proportion of 10A cells in the spheroids increases (**Fig 4(f-g))**. Similarly, as the proportion of DCIS cells in the spheroid is increased, 10A cells and their nuclei become more elongated. Thus, the presence of a more fluid-like cell type can change the shape properties of otherwise solid-like cells.

**FIG 4.**
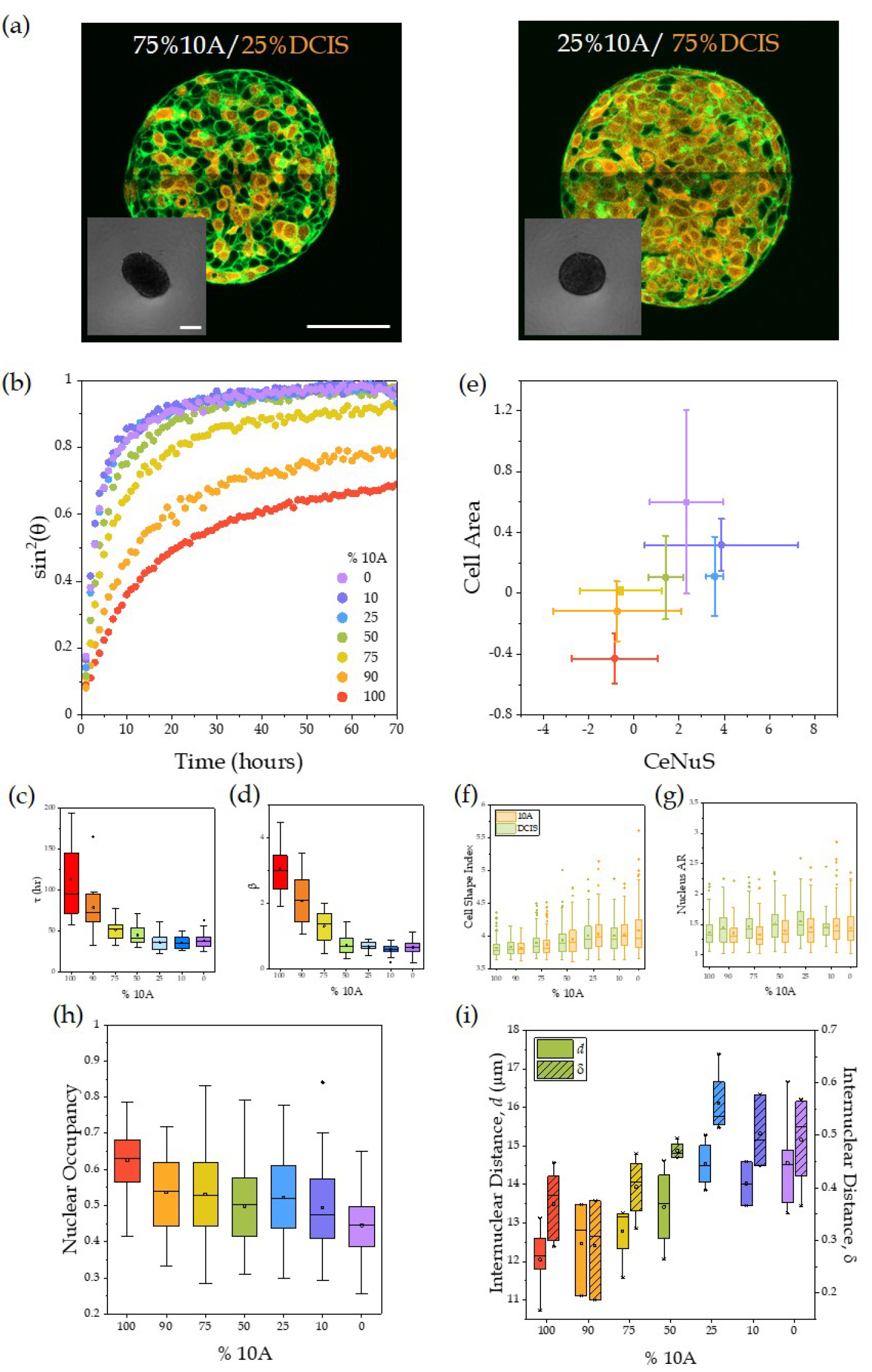
(a) Individual heterotypic 10A and DCIS spheroids with actin (green) and CellTracker-Orange labeled DCIS (orange) cells. Left spheroid contains 75% 10A cells and 25% DCIS cells, right spheroid contains 25% 10A cells and 75% DCIS cells. Scale bar is 100μm. Inset displays the corresponding fusion of heterotypic 10A/DCIS spheroids after 50 hours of fusion. Inset scale bar is 200μm. As the proportion of DCIS cells increases, fusion increases. (b) With increasing DCIS cells, fusion quickens and progresses further as demonstrated by (c) τ and (d) β values (N ≥ 15 spheroids). (e) Heterotypic 10A/DCIS spheroids in the cell area vs CeNuS phase space. There is a steady increase in both parameters as the fraction of DCIS cells increase (N ≥ 3 spheroids). Error bars display standard deviations. (f) Cell shape of individual cell types within heterotypic spheroids. Green and orange correspond to 10A and DCIS cells, respectively. (g) Nuclear shapes within heterotypic spheroids for each cell type (N ≥ 20 cells). Same legend applies to (f) and (g). (h) The nuclear-to-cell area fraction steadily decreases as more DCIS cells are added to the spheroids (N ≥ 40 cells). (i) Both absolute, *d*, and normalized internuclear, *δ* distances increase as DCIS cells are added to the spheroid (N ≥ 3 spheroids). For all box and whisker plots, whiskers display range of data excluding outliers.

The presence of the DCIS cells also loosens the packing of the nuclei, reflected in the increase of the nuclear spacing and decrease in the nuclear area fraction when DCIS cells are introduced into 10A spheroids (**Fig 4(h-i)**). Additionally, as nuclear aspect ratios increase with increasing proportion of DCIS, both the resulting absolute and normalized internuclear distances reflect the same increase in spacing between nuclei as DCIS cells are added to the system (**Fig 4(i)**). Once the effective nuclear spacing exceeds *λ* = 0.45 nuclear diameters, which occurs when 50% of the spheroid is comprised of DCIS cells, the system exhibits fluid-like behavior. In summary, we observe that in 10A-DCIS heterotypic spheroids, the DCIS-mediated fluidization process involves elongation of cells and nuclei as well as a reduction in nuclear packing that occurs through both cell volume increases and nuclear elongation.

#### 2. Heterotypic 10A-TSA spheroids

To further investigate the CeNuS vs. cell area phase space, we attempted to traverse just one axis of the phase space to examine whether the jammed 10A system could be fluidized without altering cell volumes and solely with changes to nuclear shapes. For this, we employed heterotypic spheroids composed of 10A cells and TSA-treated 10A cells (**Fig 5(a)**). As TSA-treated 10A cells are added to 10A spheroids, an increase in fusion is observed similar to that seen in 10A-DCIS mixed spheroids (**Fig 5(b)**). The corresponding τ and β values from fits decrease gradually as the proportion of TSA-treated cells increases (**Fig 5(c-d)**), reflecting the quicker and more complete fusion. Like 10A-DCIS heterotypic spheroids, 10A and 10A-TSA heterotypic spheroids exhibit dramatic changes in fluidity with only a small proportion of the fluid cell type in the system. When examining 10A and TSA-treated 10A heterotypic spheroids, the cell area-CeNuS phase space is traversed almost solely along the CeNuS axis (**Fig 5(e)**). We observe an elongation of all cells as TSA-treated cells are added. Looking at the individual cell types, cell shapes again change in the presence of another cell type (**Fig 5(f)**), while nuclear aspect ratios increase steadily with increasing TSA-treated 10A cells (**Fig 5g**). Little change is observed in the cell areas as TSA-treated cells are added.

**FIG 5.**
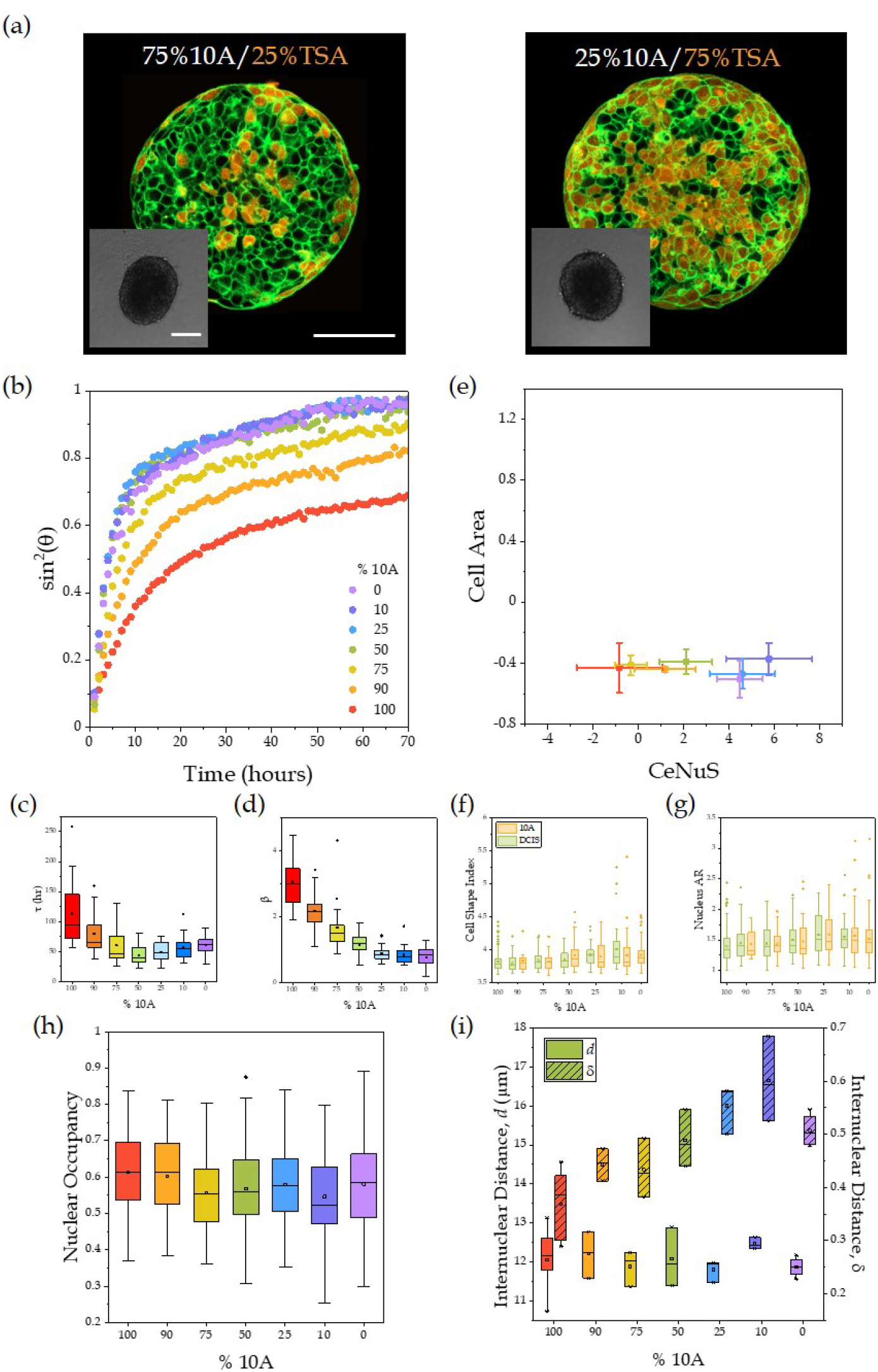
(a) Individual heterotypic 10A and TSA-treated 10A spheroids with actin (green) and CellTracker-Orange labeled DCIS (orange) cells. Left spheroid contains 75% 10A cells and 25% TSA cells; right spheroid contains 25% 10A cells and 75% TSA cells. Scale bar is 100μm. Inset displays the corresponding fusion of heterotypic 10A/TSA-treated 10A spheroids after 50 hours of fusion. Inset scale bar is 200μm. As the proportion of TSA-treated 10A cells increases, fusion increases. (b) Fusion ability increases as more TSA cells are added to 10A/TSA heterotypic spheroids. (c,d) Adding TSA cells to 10A spheroids quickens and progresses fusion (N ≥ 16 spheroids). (e) Plotted in the cell area vs CeNuS phase space, heterotypic 10A/TSA spheroids change little in cell area but increase steadily in shape index (N ≥ 3 spheroids, except for 90% 10A condition with 2 spheroids). Error bars display standard deviations. (f) Cell shape changes for individual cell types in heterotypic spheroids. Green and orange coloring represents 10A and 10A-TSA cells, respectively. Little change in cell shape is observed until over 50% of the spheroid is comprised of TSA cells. (g) Nuclear shapes elongate as more TSA cells are added to the spheroids (N ≥ 30 cells). Same legend applies to (f) and (g). (h) Little change is observed in nuclear-to-cell area fractions (N ≥ 40 cells). (i) Though there is no change in absolute distance between nuclei, once the nuclear diameter, which depends on aspect ratio, is used to normalize the distances, an increase in internuclear distance in heterotypic spheroids is apparent (N ≥ 3 spheroids). For all box and whisker plots, whiskers display range of data excluding outliers.

Along with little change in cell area, nuclear areas do not change with increasing proportions of TSA-treated cells, as exhibited by the constant nuclear-to-cell area fraction and resulting absolute internuclear distance (**Fig 5(h-i)**). Instead, nuclei become more elongated in spheroids with increasing proportion of TSA-treated cells. As a result, though no change in the absolute internuclear distances is observed, an increase in the effective nuclear spacing as TSA-treated 10A cells are added is evident (**Fig 5(i)**). Again, as in the 10A-DCIS heterotypic spheroids, fluid-like fusion occurs when the normalized internuclear distance is greater than *λ* = 0.45 nuclear diameters. However, unlike 10A-DCIS spheroids, the increase in effective nuclear spacing is due entirely to the elongated nuclei of TSA-treated 10A cells. This again supports the idea that even if nuclei are tightly packed, the addition of cells with softer nuclei can cause a shape driven increase the fusion ability of the system.

Examining all conditions on the Cell Area vs CeNuS phase space together, it is apparent that the fluid or solid-like state of the spheroids is primarily dictated by the shapes of the cells and their nuclei (**Fig 6(a)**). Specifically, fluid like behavior is reflected in elongated cells and nuclei, which translate to high CeNuS values. However, cell area also plays a role in fluidity, as for a given CeNuS value, an increase in cell area leads to an increase in fusion ability. Because nuclear area and shape are correlated with cell area and shape, we can use the single metric of normalized internuclear spacing to describe the systems studied here (**Fig 6(b)**). As internuclear distance decreases, we observe a decrease in fusion ability, with the Pearson’s correlation coefficient between *δ* and β of −0.75. Additionally, we observe a demarcation around *δ* = 0.45, below which the spheroids exhibit solid behavior and above which the spheroids fuse completely. Furthermore, by scaling nuclear area occupancy by the aspect ratio of the nuclei, another predictor of fluidity is obtained, without the need for laborious internuclear distance calculations (**Fig 6(c))**. In sum, we show that the system under study is dominated by shape-reflected and/or shape-driven changes in fusion ability. Elongated cells are better able to navigate a dense environment, not simply because of their elongated cell bodies, but also in part due to the effective increase in nuclear spacing that results from their elongated nuclei.

**FIG 6.**
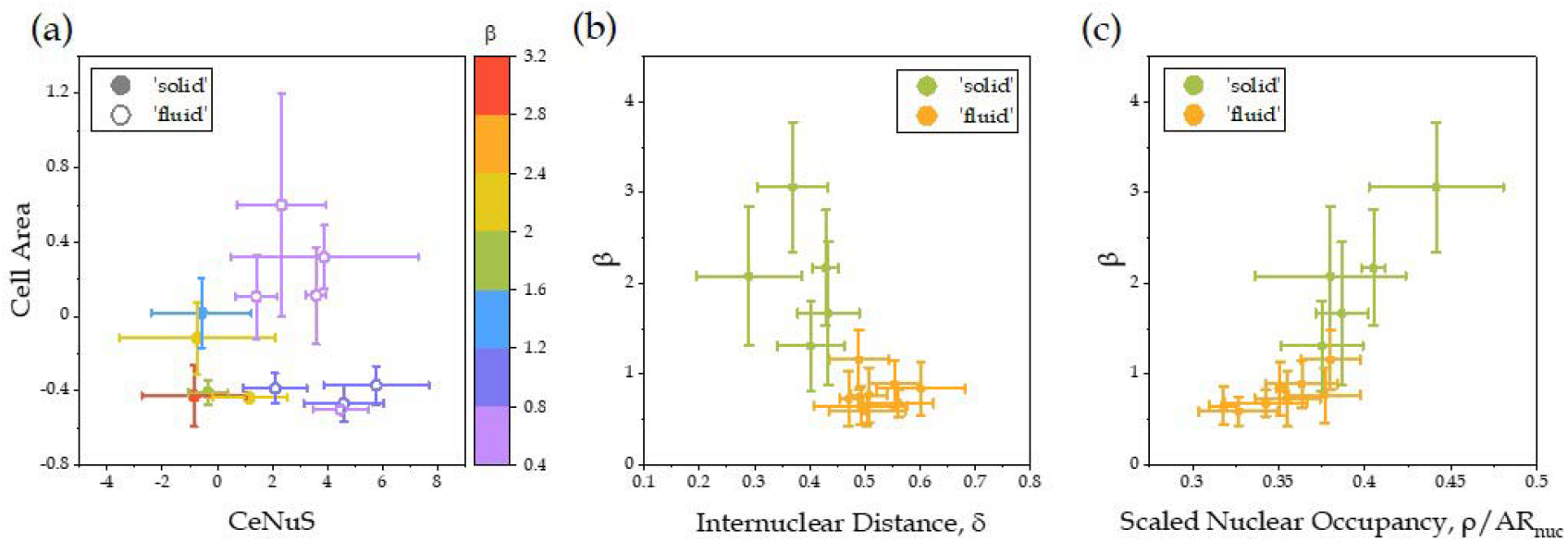
(a) All conditions displayed on the cell area vs CeNuS phase space. Coloring indicates β values and solid and open points indicate solid and fluid-like behavior, respectively. (b) The system is more fused as normalized internuclear distances, *δ*, increases (N ≥ 3 spheroids). (c) Scaled nuclear area occupancy also reflects fusion ability well, with lower occupancy correlating with more fusion (N ≥ 40 cells). Error bars display standard deviations.

## III. DISCUSSION

The results presented here agree with previous conjectures regarding the constraining effect of stiff nuclei in cell migration, and to our knowledge are the first experiments probing nuclear stiffness in multicellular spheroids. These results reveal that differences in fluidity emerge in part because of stiff and large nuclei acting as bottlenecks to cellular rearrangements in a densely packed tissue. Thus, by softening the nucleus, fluidization of an otherwise jammed system occurs. Nuclear softening leads to more deformable nuclei and subsequent changes to nuclear shape, which appear to be concurrent with changes to cell shapes. Therefore, both nuclear and cell shape serve as valuable readouts of the mechanical attributes of the cell and tissue.

In prior studies, nuclear size and occupancy (the fraction of space occupied by the nucleus within the cell) has been used as a proxy for nuclear packing density, with the underlying idea that higher nuclear occupancy leads to less intercellular space and thus smaller internuclear distances [10]. In this way, the arrangement of nuclei is essentially viewed as a packing of hard spheres [20]. We have demonstrated that nuclear occupancy in our system does in fact strongly correlate with absolute internuclear spacing. However, nuclei are deformable, and the ability of a nucleus to squeeze through internuclear spaces depends not only on absolute internuclear distances, but also on the shape that it assumes. A deformable nucleus can adopt ellipsoidal geometries with increasing aspect ratio, where the length of the minor axis becomes the limiting distance that must be traversed during cell rearrangements. By scaling internuclear distance according to this minor axis, we see that an effective increase in internuclear spacing can be achieved without increasing absolute internuclear distance, with a clear threshold denoting a regime of solid-like and fluid-like behavior. Given the correlation between nuclear occupancy and packing density, we devise a coarse-grained metric that scales the nuclear occupancy by the ellipsoidal minor axis, as a simple but accurate measurement of the effective packing density. Nuclear morphology has long played a role in cancer grading, with recent findings linking nuclear heterogeneity to higher grading [43–45]. Further, nuclear deformability and dispersion has been shown to be important to packing during germband extension, and increased nuclear occupancy has been tied to more stable isotropic packing during development [19,20,23]. Thus, our proposed metric provides another easily accessible prognostic tool that captures both shape and packing. This metric may also act as a proxy for packing efficiency, with tighter packing nuclei corresponding to more isotropic stable systems.

The MCF10A and MCF10DCIS.com system explored here offers a useful model for probing the early stages of cancer progression by capturing the transition from a normal epithelial tissue to a pre-invasive state. Surprisingly, we find that small changes in EMT status [33] lead to dramatic changes in tissue fluidity. Given that DCIS cells are generally non-invasive, fluidity is apparently acquired before invasive capacity. Additionally, a notable feature of DCIS cells is the increase in individual cell volumes relative to 10A. Recent work has pointed to volume expansion as a possible key step in advance of cancer invasion [56,57]. Volume expansion may provide a pathway for tissues to generate large forces that weaken the surrounding basement membrane, allowing for more efficient cell invasion. Such volume expansion may simultaneously fluidize the tissue through loosening of tightly packed stiff nuclei, readying its cells for invasion.

## IV. Methods and Materials

### A. Cell lines and reagents

MCF-10A cells were a gift from Professor Carol Prives (Columbia University, NY). MCF10DCIS.com were obtained from Wayne State University (Michigan, USA).

Ultralow attachment 96-well U-bottom Nunclon Sphera plates and methanol-free formaldehyde are from Thermo Fisher Scientific (Waltham, MA). Glass bottom 35mm culture dishes are from MatTek Dishes (Ashland, MA). Acetic acid (99.7%), Bovine Serum Albumine (BSA), Trichostatin A (TSA), Glycerol, Triton-X 100, hydrocortisone, cholera toxin, epidermal growth factor (EGF), insulin (10 mg/mL), ATTO-488 phalloidin, and DAPI were obtained from Sigma Aldrich (St. Louis, MO). Acid-solubilized (AS) rat tail collagen I (~4 mg/mL solution) was obtained from Advanced BioMatrix (San Diego, CA). Phosphate buffered saline (PBS) was obtained from GE Healthcare HyClone (Logan, Utah). Accutase and 100x penicillin-streptomycin-amphotericin B were obtained from MP Biomedicals (Santa Ana, CA). Horse serum and DMEM/F12 were obtained from Gibco (Grand Island, NY). CellTracker Green CMFDA and CellTracker Orange CMRA were obtained from Invitrogen Life Technologies (Grand Island, NY).

### B. Cell culture

MCF-10A and MCF10DCIS.com cells were cultured in DMEM/F12 growth medium containing 5% horse serum, 1% 100x penicillin-streptomycin-amphotericin B, 0.5μg/mL hydrocortisone, 10μg/mL insulin, 0.1 μg/mL cholera toxin, and 20ng/mL EGF. Cells were maintained at 37°C with 5% CO_2_. Cells were detached at 80-90% confluence using Accutase prior to experiments and were not used beyond passage 40.

### C. Spheroid preparation

Cells were pre-treated with 4μg/mL mitomycin-C for 2 hours to inhibit proliferation. To soften the nucleus, cells were treated with 500ng/mL Trichostatin-A (TSA) for 20 hours. We chose a TSA concentration that matches the proliferation inhibition effects of mitomycin-C.

For spheroid formation, 2000 cells were placed in wells of an ultra-low adhesion 96-well plate in media and centrifuged for 10 minutes at 1000 x g. The cells were allowed to compact for 24 hours at 37°C with 5% CO_2_ to form spheroids.

### D. Fusion microscopy

Two spheroids were placed into media-filled wells of an ultra-low adhesion 96-well plate. They were imaged every hour on a BioSpa Citation Imager using a 4x phase objective (NA 0.13). The sample was kept at 37°C and 5% CO_2_ for the duration of the experiment (up to 72 hours).

### E. Fusion analysis

The movies were thresholded and a mask was created in ImageJ. A custom Python script was written to fit the fusing spheroids to two circles and the angle and aspect ratio of their fusion was recorded. The angle of fusion, *θ*, was fit using a Kelvin-Voigt model as previously described [30]:

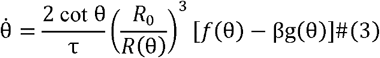

where is the *R*_0_ average initial radius of the fusing spheroids, and *R*(*θ*)is the radius of the spheroids over time. Additionally, *f* (θ)and g (θ)are defined as

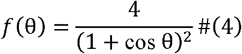

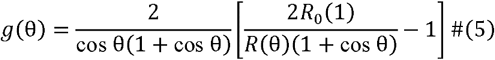

Each pair of fusing spheroids was fit using this model to obtain the resulting fit parameters, β and, *τ* in Python using SciPy’s curve_fit function.

### F. Labelling and Imaging

Low tolerance dishes were collagen coated with 50μg/mL AS collagen for 1 hour, then rinsed with PBS and water. Spheroids were fixed with 4% formaldehyde for 1 hour, washed three times for 10 minutes each with PBS, permeabilized with 0.2% Triton-X 100 for 1 hour, washed three times for 10 minutes each with 10% (w/v) bovine serum albumin (BSA) and soaked in 10% BSA for 30 minutes. Spheroids were then placed on collagen coated glass bottom dishes. The spheroids were then labeled with 1:400 ATTO 488 phalloidin and 1:500 DAPI in 10% BSA for 24 hours at 4°C. The spheroids were then washed with BSA, mounted in 80% glycerol with pH of 8.5, and left to clear for at least 24 hours at 4°C. For cells where cytosol labeling was desired, cells were incubated with CellTracker Orange for 45 minutes at a concentration of 2μM prior to spheroid formation.

Fixed spheroid imaging was performed on an inverted confocal laser-scanning microscope (Zeiss LSM800) in confocal fluorescence mode with 20x air (NA 0.75) or 63x oil (NA 1.4) objectives. Spheroids were imaged around their bisecting plane.

### G. Cell shape quantification

Cell and nuclear shapes were analyzed manually. Nuclear positions were found using the StarDist package in ImageJ [58], with adjustments made by hand to exclude small debris that was selected or to include nuclei that were missed. For each spheroid, a single CeNuS parameter (Eqn. (2)) was calculated by sampling a subset of cells within the spheroid. The values were standardized to a random subset of the mitomycin-C treated 10A and DCIS cell data.

### H. Nearest neighbor calculation

Nuclear positions were identified using the StarDist plugin in ImageJ [45]. A K-Means tree was constructed with all nuclei in the spheroid. The distances of the 6 nearest neighbors of each nucleus were calculated and averaged to obtain the average nuclear spacing for each spheroid. This quantity was then normalized by the average shortest axis of the nucleus of that spheroid type using the aspect ratio and area of the nuclei, obtained from ImageJ (Eqn. (1)). Trends were found to be invariant to the number of nearest neighbors probed.

### I. Statistical analysis

All data was tested for normality using Shapiro Wilks tests. If normal, t-tests were performed (indicated by p_t-test_). Otherwise, Mann-Whitney tests were performed (indicated by p_MW_). To test correlations, a Pearson’s correlation test was performed. All tests were performed in OriginPro. Statistical significance is indicated by *p<0.05, **p<0.01, ***p<0.001, ****p<0.0001. Lack of statistical significance is denoted by a dagger symbol (†).

## Acknowledgements

These studies used the Confocal and Specialized Microscopy Shared Resource of the Herbert Irving Comprehensive Cancer Center at Columbia University, funded in part through the NIH/NCI Cancer Center Support Grant P30CA013696.

## APPENDIX A CORRELATION BETWEEN ASPECT RATIO AND β

Linear regression was performed to assess the relationship between aspect ratio (*AR*) and β in OriginPro. The resulting fit has an adjusted *R*^2^ value of 0.92 with p-values <0.01 for both the intercept and slope: *AR =* 0.99 + 0.18 *β*.. Thus, *AR* =1.2 equated to β = 1.2.

**FIG 7.**
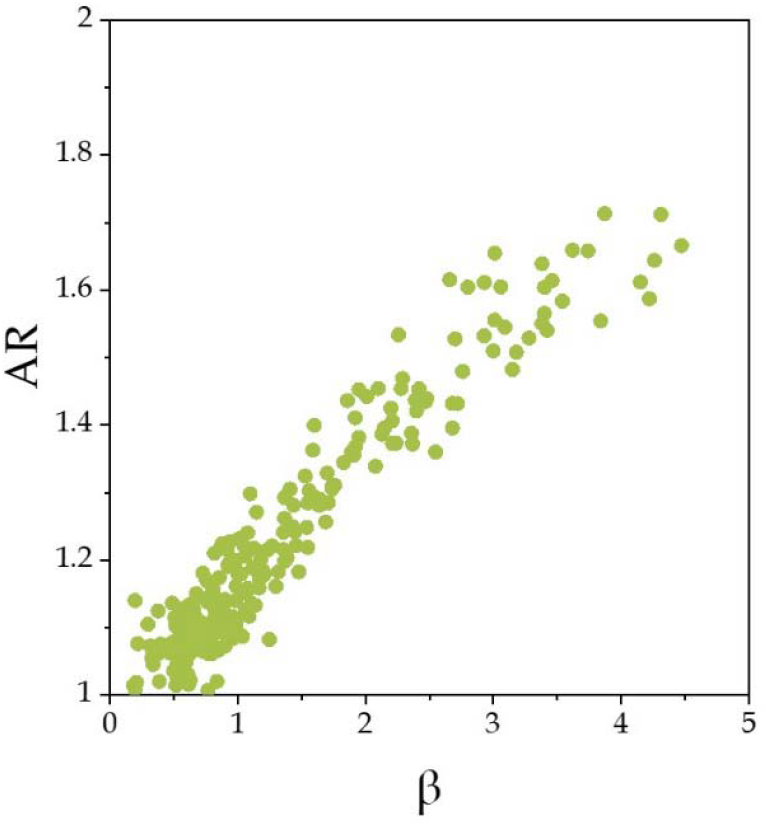
Aspect ratio plotted against β from Kelvin-Voigt fits. Dotted line displays fit from linear regression with *R*^2^ =0.92.

## APPENDIX B PROLIFERATION EFFECTS

We investigated the effects of proliferation on coalescence. For 10A spheroids, coalescence is progressed by proliferation, while for DCIS little change is observed.

**FIG 8.**
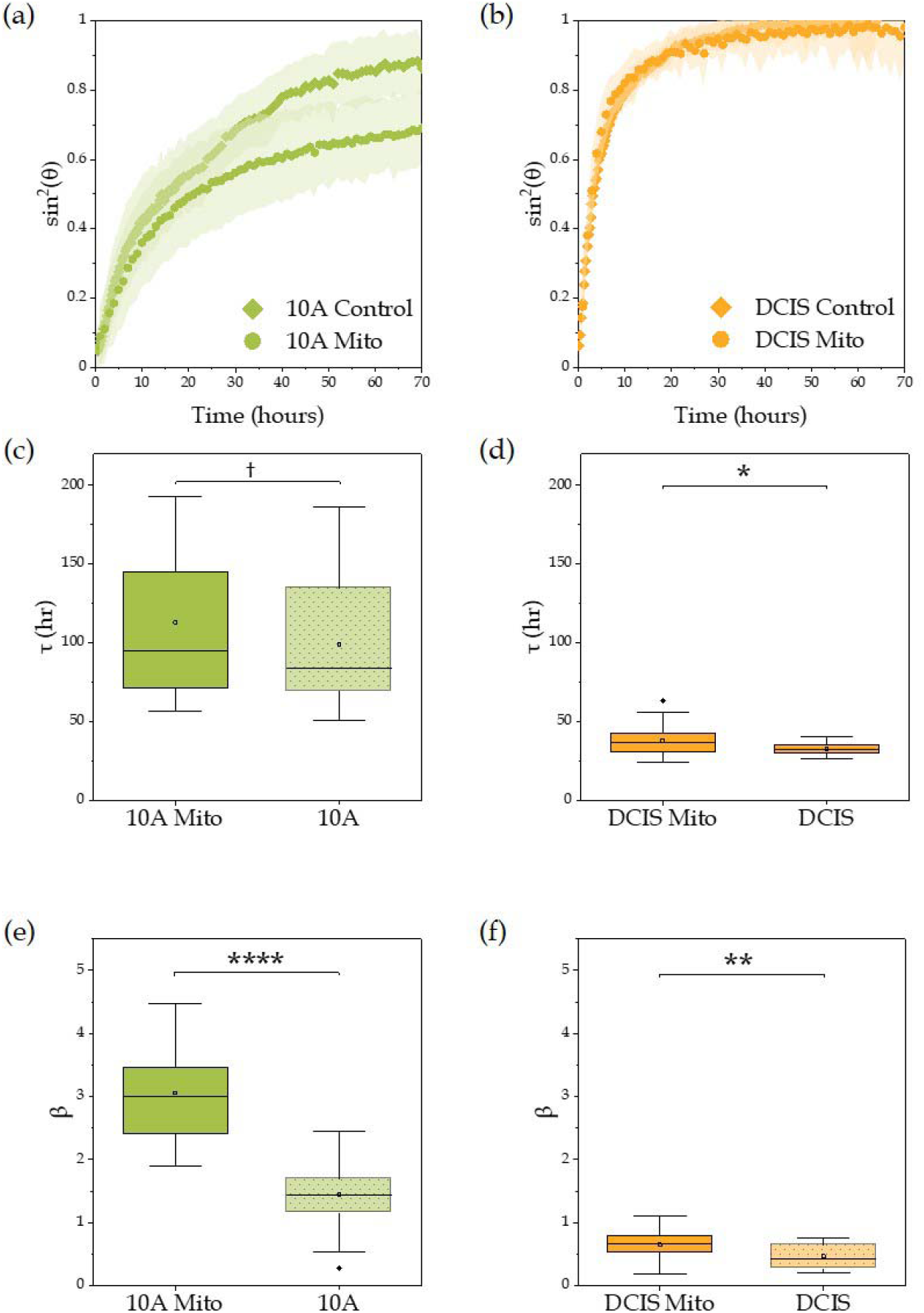
(a,b) Evolution of the angle of fusion via sin^2^θ, with θ the angle of fusion for (a) 10A and (b) DCIS spheroids treated with mitomycin-C vs. control, with the mitomycin-C treated data also shown in Fig. 1 in Main Text. (c) Box plot displaying no significant change in τ from Kelvin-Voigt fits between mitomycin-C and control 10A spheroids (p_MW_ > 0.05). (d) Control DCIS spheroids fuse faster than mitomycin-C treated DCIS spheroids (p_t-test_ < 0.05). (e,f) β from fits to the Kelvin Voight model for control and mitomycin-C treated sets of (e) 10A (p_t-test_ < 0.0001) and (f) DCIS spheroids (p_t-test_ < 0.01). (N ≥ 16 spheroids).

## APPENDIX C CELL SHAPES AT FUSION NECKS

Rapid initial fusion is due in part to the elongated cells at the edge of the spheroids. These elongated borders come into contact at the neck of fusing spheroids.

**FIG 9.**
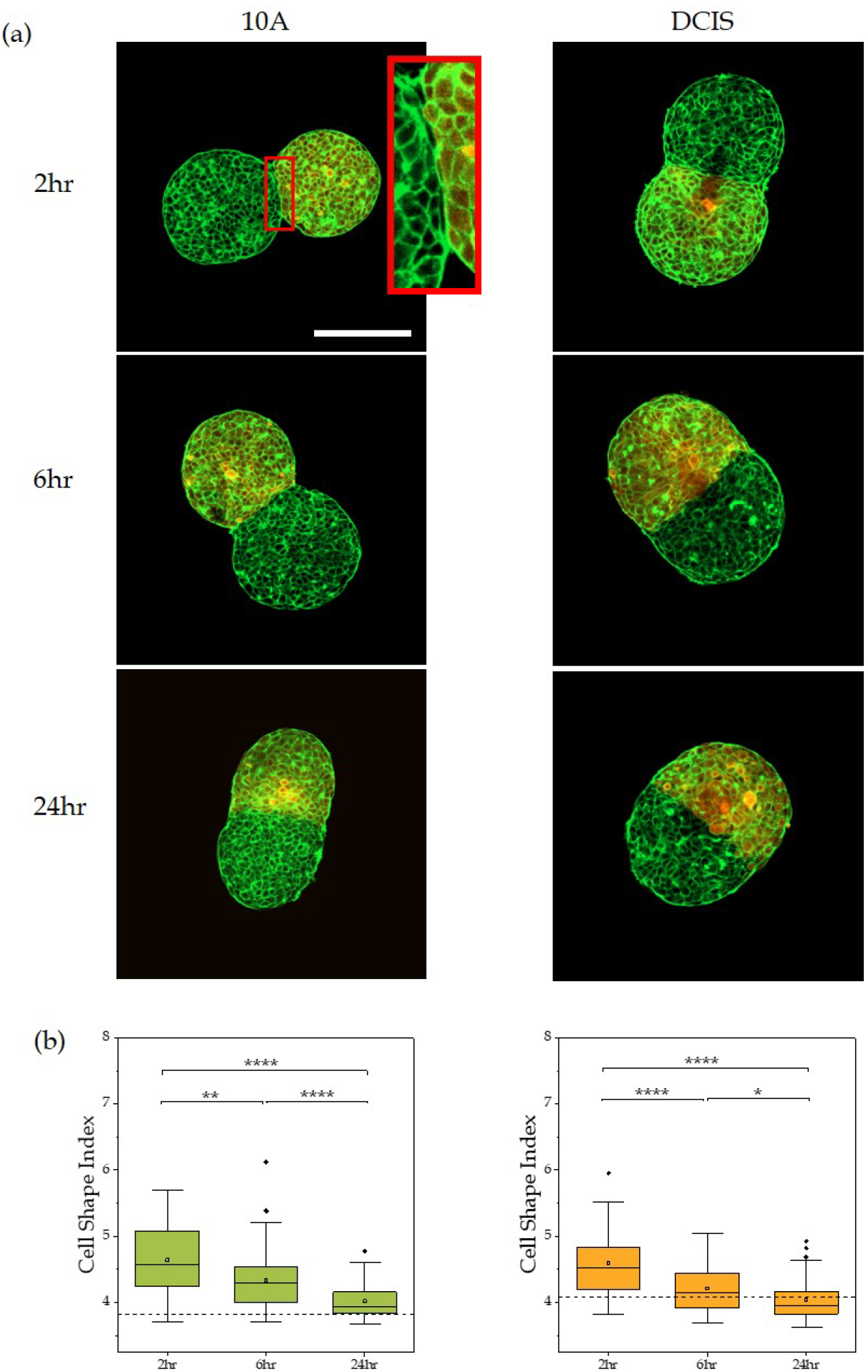
(a) Fusion progression for (left) 10A and (right) DCIS spheroids at (top) 2 hr and (bottom) 24 hr. One spheroid is labeled only with phalloidin (green) while the other is labeled with phalloidin and Cell Tracker Orange (orange). Inset displays higher magnification image of neck cells in contact with the other spheroid. Scale bars are 100 µm. (b) Shape evolution of cells at the neck of fusing spheroids over time. (n ≥ 40 cells). Dotted reference line displays the cell shape index of the bulk of the corresponding individual spheroids.

## APPENDIX D TRICHOSTATIN-A-TREATED DCIS SPHEROIDS

TSA leads to a small decrease in fusion timescale and increase in fusion for DCIS spheroids. Little change in the shapes and a small decrease in nuclear occupancy are observed.

**FIG 10.**
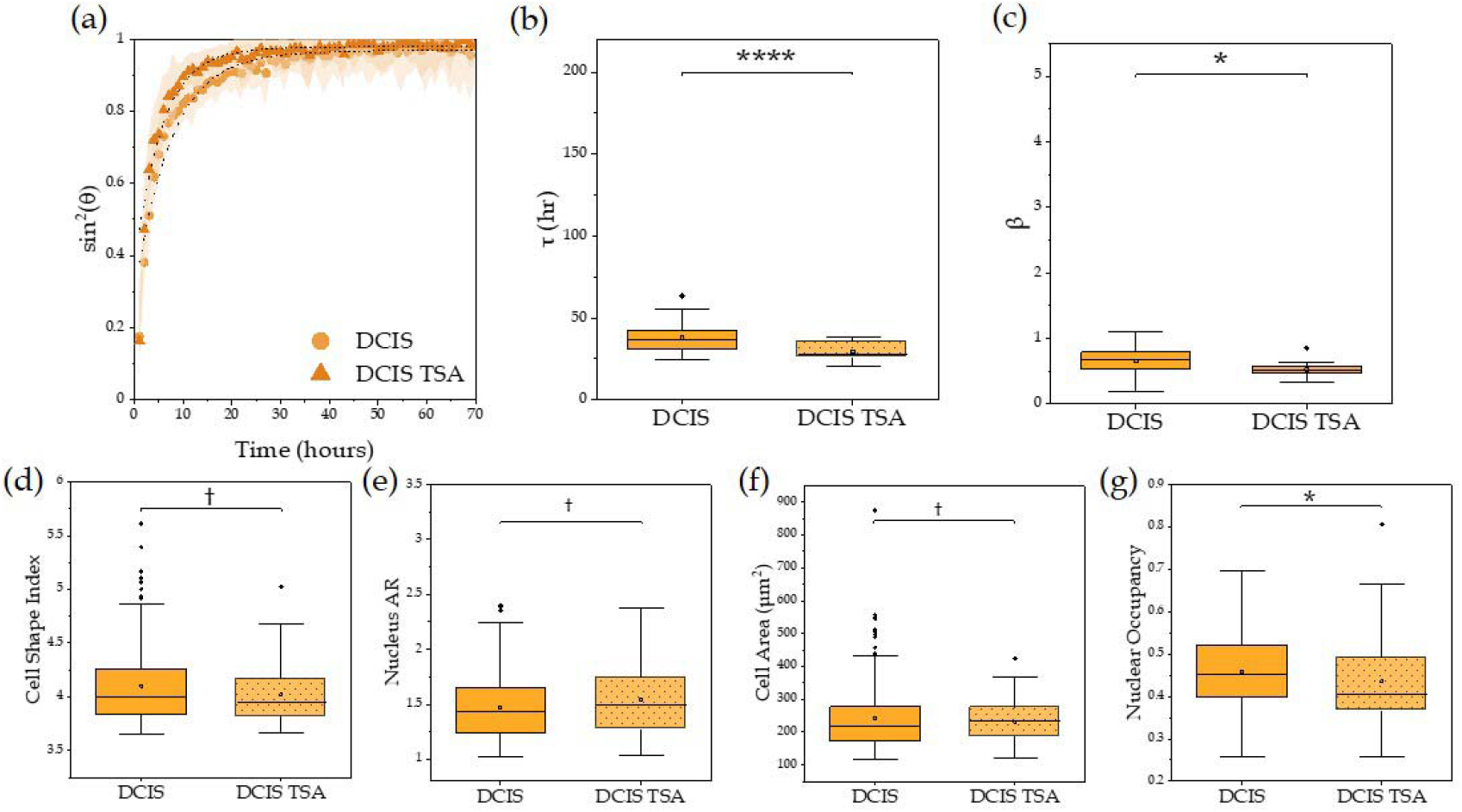
(a) TSA-treated DCIS spheroids (triangles) fuse slightly more than control spheroids. (b,c) TSA-treated DCIS spheroids fuse slightly faster and more, as demonstrated by lower (b) τ (p_t-test_ < 0.0001) and (c) β (p_t-test_ < 0.0001) values (N ≥ 16 spheroids). (d-f) TSA treatment has little effect on (d) cell shape index, (e) nuclear AR, (f) cell area (p_MW_ > 0.05), and (g) nuclear occupancy fraction (p_MW_ < 0.05) (N > 50 cells).

